# Interplay between environmental yielding and dynamic forcing regulates bacterial growth

**DOI:** 10.1101/2023.12.05.569991

**Authors:** Anna M. Hancock, Sujit S. Datta

## Abstract

Many bacterial habitats—ranging from gels and tissues in the body to cell-secreted exopolysaccharides in biofilms—are rheologically complex, undergo dynamic external forcing, and have unevenly-distributed nutrients. How do these features jointly influence how the resident cells grow and proliferate? Here, we address this question by studying the growth of *Escherichia coli* dispersed in granular hydrogel matrices with defined and highly-tunable structural and rheological properties, under different amounts of external forcing imposed by mechanical shaking, and in both aerobic and anaerobic conditions. Our experiments establish a general principle: that the balance between the yield stress of the environment that the cells inhabit *σ*_*y*_ and the external stress imposed on the environment *σ* regulates bacterial growth by modulating transport of essential nutrients to the cells. In particular, when *σ*_*y*_ *< σ*, the environment is easily fluidized and mixed over large scales, providing nutrients to the cells and sustaining complete cellular growth. By contrast, when *σ*_*y*_ *> σ*, the elasticity of the environment suppresses large-scale fluid mixing, limiting nutrient availability and arresting cellular growth. Our work thus reveals a new mechanism, beyond effects that change cellular behavior via local forcing, by which the rheology of the environment may regulate microbial physiology in diverse natural and industrial settings.

---

Many bacterial environments—e.g., gels and tissues inside hosts, subsurface soils and sediments, exopolysaccharides in biofilms and in the environment, activated sludge in sewage treatment plants, and food products [1–13]—are neither perfectly elastic solids nor simple viscous fluids. Instead, they are yield stress materials: they behave as viscoelastic solids when exposed to weak mechanical stresses, but flow when the imposed stress exceeds a threshold yield stress *σ*_*y*_. Indeed, these habitats are typically not quiescent, but are subjected to continuous and dynamic external forcing by moving boundaries, fluid flows, and other mechanical stressors. Thus, depending on the balance between the yield stress *σ*_*y*_ and external stresses *σ*, the environments that bacteria inhabit can vary between being solid-like and liquid-like.

A familiar example is mucus, which serves as a habitat for both commensal and pathogenic bacteria in diverse animals. In healthy humans, airway mucus is often a runny solution with a small or negligible yield stress; however, many respiratory disorders are characterized by more concentrated mucus whose yield stress can be as large as tens of Pascals [14–18]. As a result, the beating cilia that line the airways are less effective at clearing mucus from the lungs—leading, in some cases, to chronic and deadly infections [15, 19]. Another familiar example is the polymer matrix that encapsulates the cells in a surface-attached biofilm, such as the plaque that we brush off our teeth and the slime that can grow on industrial equipment, medical catheters, and even in our showers. In some cases, this matrix is weak and easily yielded, whereas in others, its yield stress can be as large as thousands of Pascals [13, 20– 23]—which is thought to contribute to biofilm virulence and recalcitrance to treatment. Given that the yield stress of the environments that bacteria inhabit can vary so widely (Table I), with implications for health, environment, and our everyday lives, we ask: how do changes in yield stress influence bacterial behavior?

**Table 1.** Order of magnitude estimates of rheological parameters characterizing bacterial environments in nature and in this study. Mucus yield stress values are drawn from studies with human and porcine lung and gastric mucus and biofilm EPS yield stress values are drawn from studies of *Pseudomonas aeruginosa* and *Staphylococcus epidermidis*. External stresses for mucus include examples of ciliary clearance, coughing, mechanical ventilation, and digestion. External stresses for biofilms include examples of mechanical agitation from tooth brushing, mixed bioreactors, and shear cleaning of biofouling on ship hulls.

| Environment | Yield stress $\sigma_y$ (Pa) | External stress $\sigma$ (Pa) | $Bi \equiv \sigma_y/\sigma$ | References |
| --- | --- | --- | --- | --- |
| Lung mucus: healthy | $10^{-1} - 10$ | $10^{-4} - 10^4$ | $10^{-5} - 10^5$ | [14–18, 24–29] |
| Lung mucus: cystic fibrosis | $10^{-1} - 10^2$ | $10^{-2} - 10^5$ | $10^{-6} - 10^4$ | [14–17, 24–29] |
| Biofilm EPS | $10^{-1} - 10^2$ | $10^{-2} - 10^6$ | $10^{-7} - 10^4$ | [13, 20–35] |
| Granular hydrogel matrix | $\ll 10^{-2} - 10^2$ | $0 - 1$ | $\ll 10^{-2} - \gg 10^2$ | This study, [36–38] |

Prior research has investigated how other aspects of environmental rheology can influence bacterial behavior via *local* mechanical forcing [39, 40]. For example, a growing body of work is elucidating the ways in which individual bacterial cells on 2D planar surfaces sense and respond to changes in surface stiffness and topography via local mechanical forces, using their flagella, pili, and membrane proteins [41, 42]. Other work has shown how 3D confinement of dense, multicellular colonies causes cells to rearrange, slow down growth, and even induce biofilm formation due to large cell-cell contact forces [43–48]. In bulk liquids, studies have shown how local stresses generated by fluid flows generated by bacterial swimming alter swimming kinematics [49–54].

Here, we report another, distinct mechanism by which environmental rheology impacts a fundamental aspect of bacterial physiology—cellular growth. By studying *Escherichia coli* growth inside permeable 3D granular hydrogel matrices, we show that the balance between the yield stress *σ*_*y*_ and external stress *σ*, quantified by the Bingham number Bi ≡*σ*_*y*_*/σ* and tuned over a wide range in our experiments, regulates bacterial growth by modulating transport of externally-supplied essential nutrients to the cells. In particular, when the matrix is fragile enough to be fluidized by shaking (small Bi), mixing transports oxygen from the boundaries of the matrix to the cells; as a result, the bacteria are able to perform aerobic respiration and continue through their entire growth cycle. In stark contrast, when the matrix is tough enough to withstand shaking (large Bi), its elasticity hinders mixing; consequently, the majority of the bacteria become oxygen-depleted and their growth cycle is arrested. Notably, owing to the unique structure of the hydrogel matrices, this transition between continued and arrested growth is not associated with changes in the local mechanical environment encountered by individual cells. Hence, this mechanism by which environmental rheology regulates bacterial physiology—by modulating large-scale transport of nutrients—complements other mechanisms that rely on local mechanical forcing instead. Because many bacterial habitats have complex rheological properties, encounter dynamical external forcing, and have heterogeneous nutrient distributions, we anticipate that our findings are applicable to microbial life in diverse natural and industrial settings.

## Materials and Methods

### Details of bacterial strains and growth media

Our experiments use two different strains of *E. coli*, both of strain background W3110 that constitutively express GFPmut2: a motile strain as well as a non-motile Δ*flhDC* mutant. In all experiments, *E. coli* are grown in EZ Rich, a defined, rich growth medium (Teknova; Hollister, CA). Our procedure to prepare 100 mL of liquid EZ Rich is as follows: Mix 10 mL of 10X MOPS mixture (M2101), 10 mL of 10X ACGU solution (M2103), 20 mL of 5X Supplement EZ solution (M2104), and 1 mL of 0.132 M potassium phosphate dibasic solution (M2102) in 58 mL of ultrapure milli-Q water. We then autoclave this mixture and add 1 mL of sterile 20% aqueous glucose solution once the mixture has cooled.

### Preparing granular hydrogel matrices

We use dense packings of hydrogel grains (“microgels”) as growth matrices for bacteria (Figure 1A). Each matrix is prepared by dispersing dry granules of internally cross-linked microgels made of biocompatible acrylic acid-alkyl acrylate copolymers (Carbomer 980; Lubrizol, Wickliffe, Ohio) in liquid EZ Rich. The granules absorb the liquid until their elasticity prevents further swelling. We ensure a homogeneous dispersion of swollen microgels by mixing for at least 12 h using either magnetic stirring or a rotary mixer, and adjust the final pH to 7.4 by adding 10 M NaOH. The cells are then uniformly dispersed in the liquid phase in between the microgels and their growth monitored using spectroscopy, as described further below.

**FIG. 1.**
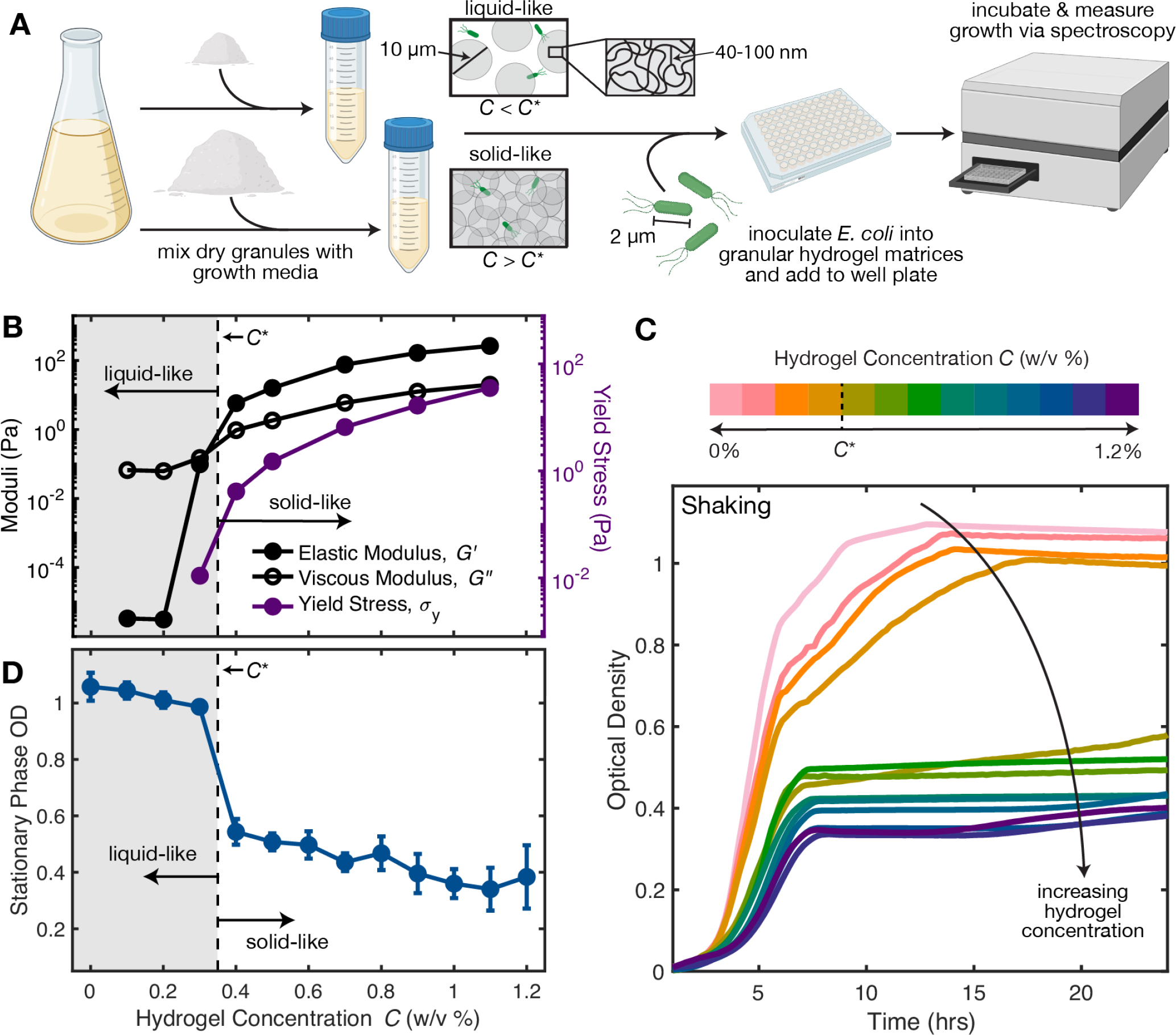
Bacterial growth is tuned by environmental rheology. **(A)** Schematic showing an overview of the experiments. (Left) We prepare granular hydrogel matrices by mixing dry hydrogel granules with liquid growth media at prescribed mass fractions *C*. (Middle) The individual granules swell to form permeable microgels whose internal mesh permits growth substrates to freely diffuse, but acts as a solid barrier that confines cells. Tuning *C*, and therefore how close-packed the swollen microgels are, enables us to tune the overall rheological properties of the hydrogel matrices. (Right) In each hydrogel matrix, we disperse a dilute inoculum of *E. coli* in the interstitial voids between microgels and use spectroscopy to measure subsequent bacterial growth over 24 h under shaken or non-shaken conditions. As shown by the shear rheology measurements in **(B)**, for small *C < C*^*\**^, the dilute suspension of microgels is liquid-like; the yield stress *σ*_*y*_ (purple symbols) is below the measurement resolution, and the viscous loss modulus *G*^*″*^ (open black symbols) exceeds the elastic storage modulus *G*^*′*^ (filled black symbols). By contrast, when *C > C*^*\**^, the microgels are jammed, and the packing is solid-like; *σ*_*y*_, *G*^*′*^, and *G*^*″*^ all precipitously increase as *C* increases above the jamming concentration *C*^*\**^, and *G*^*′*^ *> G*^*″*^. **(C)** *E. coli* growth curves measured using spectrophotometry under shaken conditions in granular hydrogel matrices with varying *C*. Color bar denotes the hydrogel concentration where each color change corresponds to a 0.1% change in *C*. Each curve is an average of 3-5 replicates from 1 experiment. The bacteria follow a complete growth curve for *C < C*^*\**^ (pink to orange), but growth is arrested when *C > C*^*\**^ (brown to purple). **(D)** The transition between complete and arrested growth is summarized by plotting the stationary phase optical density, defined as the average OD_600_from *t* = 18*−*24h of the growth curves, as a function of *C*. The stationary phase OD_600_ drops abruptly as *C* increases above *C*^*\**^. Error bars represent standard deviation from replicates across *>* 3 independent experiments.

The swollen microgels are *∼*5-10 µm in diameter with *∼*20% polydispersity, but have an internal mesh size of *∼*40–100 nm, which permits growth substrates such as amino acids, glucose, and oxygen to freely diffuse throughout while impeding cellular motion [37]. Our experiments use matrices of different mass fractions of dry granules, *C* (w/v %), resulting in different packing fractions of the swollen microgels—ranging from dilute suspensions with a small *σ*_*y*_ as described further below, to tightly-jammed packings with a large *σ*_*y*_ (schematized in the middle of Fig. 1A). Hence, our approach yields hydrogel matrices whose microstructure does not limit the ability of cells to access small molecules, but whose macroscopic rheological properties can be systematically tuned to mimic bacterial environments.

### Rheology of granular hydrogel matrices

A unique feature of our hydrogel matrices is that their rheological characteristics can be dramatically altered by a small change in microgel concentration [37, 38]—enabling us to probe the limits of small and large *σ*_*y*_ by varying the mass fraction *C*. We quantify these characteristics with an Anton Paar (Graz, Austria) MCR501 shear rheometer, loading ~2 mL of each dispersion in the 1 mm gap between two roughened 50 mm-diameter parallel circular plates. In particular, for each matrix, we use unidirectional steady shear rheology to measure the variation of the shear stress *σ* with shear rate 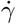 (Fig. S1). For sufficiently large *σ > σ*_*y*_, each matrix is fluidized and *σ* increases with increasing 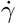 ; extrapolating to 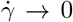 by averaging the measured shear stress for 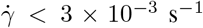 then provides an estimate for *σ*_*y*_. Our measurements are summarized by the purple circles in Fig. 1B. Matrices with small *C* (indicated by the grey shading) are liquid-like; their yield stress is below or near the rheometer measurement resolution, characteristic of a dilute suspension of swollen microgels. By contrast, just above *C* = *C*^*\**^ ≈0.35%, the matrices become solid-like, and *σ*_*y*_ increases precipitously—indicating that the microgels are sufficiently dense-packed to form a jammed packing. We identify the transition between these two rheological regimes as the jamming concentration, *C*^*\**^, shown by the dashed line in Fig. 1B [37, 38]. As *C* continues to increase above *C*^*\**^, *σ*_*y*_ increases as is characteristic of jammed packings. The values of *σ* range from *∼*10^*−*2^ to *∼*10^2^ Pa — spanning the range of yield stresses encountered by bacteria in many natural habitats (Table I)

As a further corroboration of this rheological jamming transition, we use oscillatory rheology to measure the linear elastic storage and viscous loss moduli *G*^*′*^ and *G*^*″*^, respectively, of each matrix (Fig. S2). The variation of both moduli at a fixed strain amplitude of 1% and a fixed oscillation frequency of 1 Hz with *C* is shown by the black circles in Fig. 1B; these values are chosen so that the measurements are independent of strain amplitude (Fig. S2A) and oscillation frequency (Fig. S2B). Consistent with our expectation [37, 38], *G*^*″*^ (open circles) exceeds *G*^*′*^ (filled circles) for matrices with *C < C*^*\**^, indicating that viscous dissipation dominates over the storage of elastic stresses—that is, these matrices are liquid-like. By contrast, for matrices with *C*≥*C*^*\**^, *G*^*′*^ exceeds *G*^*″*^, indicating that the network of inter-grain contacts and deformations of individual grains can support the applied stress and dominates the rheological response—these matrices are solid-like.

### Measuring aerobic bacterial growth

How do bacteria grow within matrices of different *σ*_*y*_? Notably, due to the high degree of microgel swelling and small polymer concentration (*C*≤1.2%), our granular hydrogel matrices are transparent. Therefore, we directly probe the growth and proliferation— hereafter referred to as “growth” for brevity—of *E. coli* using UV-Vis spectroscopy (Biotek Epoch 2 microplate spectrophotometer; Winooski, Vermont), starting with a dilute suspension of cells uniformly dispersed between the microgels (Fig. 1A).

To achieve this starting condition, prior to each experiment, we pick a single colony from an agar plate and grow it in liquid EZ Rich for 24 h in a 37^*°*^C shaking incubator. The next day, we dilute the overnight culture to an optical density (OD) of *∼*1.5. We then inoculate 3*−*5 mL of each test granular hydrogel matrix in a conical tube with 30-50 µL of the diluted overnight culture and uniformly mix the cells throughout using vigorous shaking, marking *t* = 0h for the subsequent growth curves. We remove the bubbles introduced by mixing by centrifuging at 3000 RCF for 10 s. Then, we gently deposit 200µL of each inoculated matrix into the cylindrical wells of a 96-well cell culture plate (Corning Incorporated, Kennebuck, Maine) using a pipette or 1 mL syringe for liquidlike or solid-like matrices, respectively. We also aliquot sterile replicates of each hydrogel matrix without *E. coli* to use as a “blank” reference OD_600_ for each *C*.

We then characterize the full bacterial growth curve by incubating the microplate at 37^*°*^C for 24 h, measuring UV-vis absorption at 600 nm (OD_600_) every 10–15 min. To test the influence of dynamic forcing on growth, some experiments are performed shaken, in which the microplate undergoes continuous linear shaking at 567 cycles per min over a distance of 3 mm, while others are non-shaken, static cultures. We represent the growth curves (Figs. 1C, 2A,C) by plotting the blank-subtracted OD_600_ over time. The stationary phase OD_600_ of each sample is given by the average value of the blank-subtracted OD_600_ 18-24 h after inoculation.

To validate the stationary OD_600_ measurements obtained using the aerobic microplate reader, we also measure the number density of colony forming units (CFUs/mL) for stationary phase cell cultures in granular hydrogel matrices of different concentrations. To do so, we first grow *E. coli* for 24 h in the matrices in microplate wells under both shaken and non-shaken conditions, as described above. We then extract ≈20-80 µL from the center of each microplate well for serial dilution and plating, and verify the pipetted volumes of these samples using mass measurements (OHAUS PA84C, Parsippany, New Jersey) assuming a density of *≈* 1 g*/*mL.

Next, we serially dilute these samples by factors of 10 in phosphate-buffered saline, and plate three 10 µL droplets of each dilution on plates made of 2% Lennox LB Broth (Sigma Aldrich, St. Louis, Missouri) and 1.5% agar (Sigma Aldrich, St. Louis, Missouri). We incubate the plates at 30^*°*^C overnight before counting individual colonies the following morning. The results thereby obtained (Fig. S3) show good agreement with the results obtained using OD_600_ measurements (Fig. 2B); in both cases, we observe a discontinuous transition in growth between two states for shaken cultures of increasing hydrogel concentration *C*, whereas growth is not sensitive to *C* for non-shaken cultures.

**FIG. 2.**
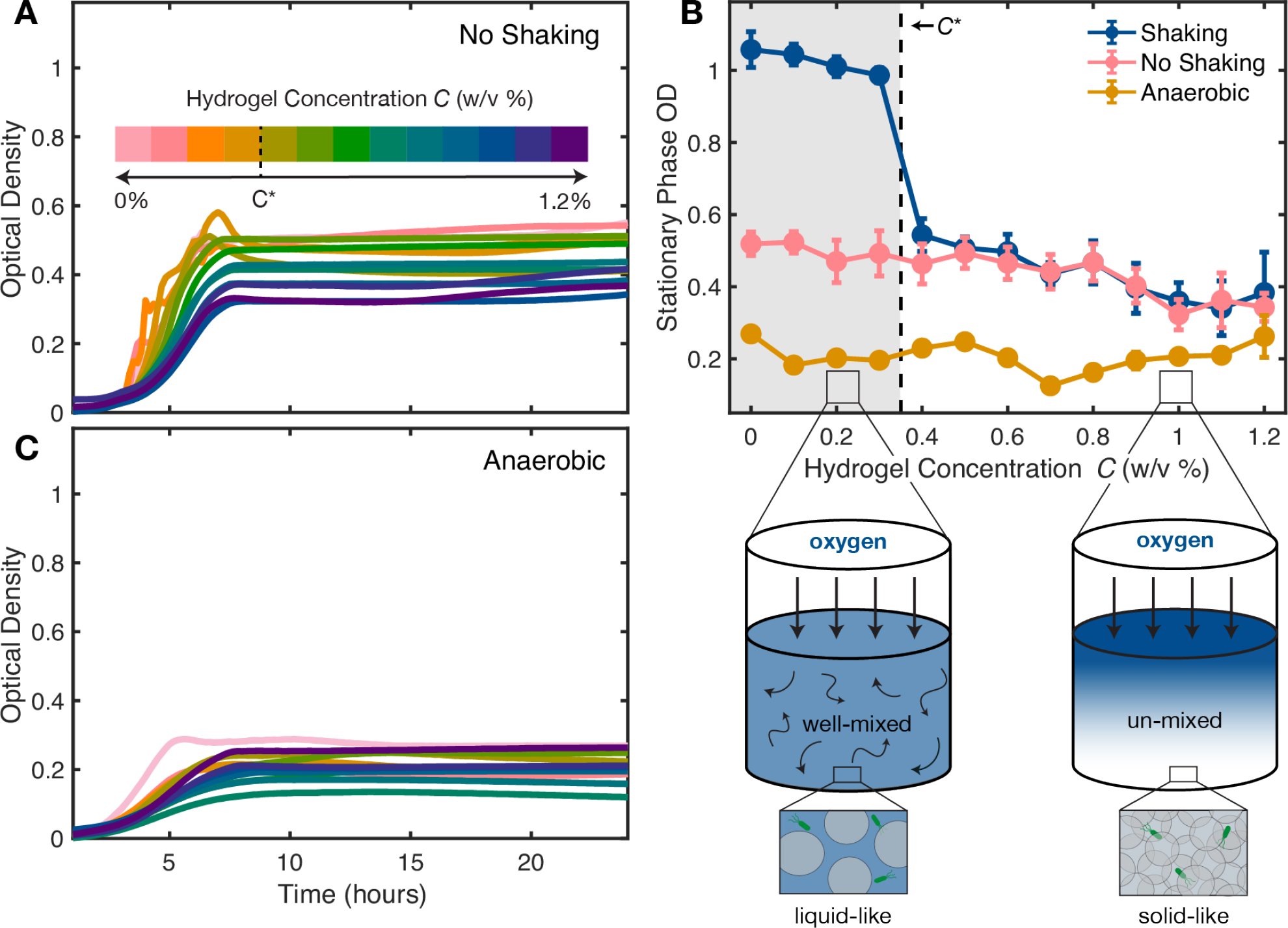
Bacterial growth is arrested across all hydrogel concentrations in non-shaken aerobic conditions and in shaken anaerobic conditions. **(A)** *E. coli* growth curves measured under non-shaken conditions in hydrogel matrices with varying *C*; in this case, cells exhibit arrested growth across all hydrogel concentrations (compare to Fig. 1C). **(C)** *E. coli* growth curves measured under shaken but anaerobic conditions in hydrogel matrices with varying *C*; cells again exhibit arrested growth across all hydrogel concentrations. Each curve in **(A)** and **(C)** is an average of 3-5 replicates from 1 experiment. **(B)** These measurements are again summarized by plotting the stationary phase OD_600_ as a function of *C*. While cells in shaken aerobic conditions exhibit a transition from complete to arrested growth across the jamming concentration (blue points, Fig. 1), cells in non-shaken aerobic conditions (pink) or shaken anaerobic conditions (blue) show arrested growth across all *C*. Error bars represent standard deviation from replicates across 2-6 independent experiments. These measurements indicate that in aerobic conditions, when shaking is sufficient to fluidize the hydrogel matrix (*C < C*^*\**^), mixing provides externally-supplied oxygen to the cells and sustains complete growth (left schematic)—while in non-shaken conditions, oxygen transport is diffusion-limited and therefore cellular growth is arrested (right schematic).

### Measuring anaerobic bacterial growth

To test the influence of oxygen availability on bacterial growth within the granular hydrogel matrices, we also measure growth curves under anaerobic conditions. To do so, we use a Biotek 800 TS Absorbance Microplate Reader (Winooski, Vermont) contained within a temperature-controlled Coy Laboratory Products (Grass Lake, Michigan) vinyl anaerobic chamber that maintains oxygen concentrations below 60 ppm, which corresponds to *∼*0.03% of atmospheric oxygen at sea level. The experimental procedure, including shaking settings, is identical to that for measuring aerobic growth with one notable exception: prior to inoculation by the bacteria—which are grown aerobically overnight—each granular matrix is purged of oxygen in the anaerobic chamber for 24 h. All subsequent steps, including inoculation and microplate preparation, are performed in the anaerobic chamber. Additionally, the anaerobic Absorbance Reader reads OD at 630 nm instead of 600 nm. As shown in Fig. S4, OD measurements performed on the same liquid bacterial cultures using both aerobic and anaerobic microplate readers are linearly proportional; we therefore use this proportionality to convert the anaerobic OD_630_ measurements to OD_600_.

### Probing oxygen limitations using confocal microscopy

It is well established that newly synthesized GFP requires *>* 0.06% of molecular oxygen to complete fluorophore formation and emit an appreciable fluorescent signal [55–57]. We use this oxygen dependence to probe the spatial distribution of oxygen in the different experiments. To do so, we repeat both shaking and non-shaking growth experiments with glass-bottom 96-well plates (Cellvis, Sunnyvale, CA) for non-motile *E. coli* constitutively expressing GFPmut2 in *C* = 0.2% and 0.6%. For shaking experiments, plates were incubated in the Biotek Epoch 2 Absorbance Reader at 37^*°*^C with identical shaking conditions to growth experiments. For non-shaking experiments, cells were incubated in a stationary incubator at 37^*°*^C. After ≈20 hours of growth, we remove plates from their incubation condition and use a Nikon A1R+ inverted laser-scanning confocal microscope to acquire vertical stacks of planar fluorescence and brightfield images, each separated by 2.5 µm in depth, near the bottom surface of each well. We then report the average GFP fluorescence intensity of the planar image 15µm above the bottom surface for each replicate in Fig. 3.

**FIG. 3.**
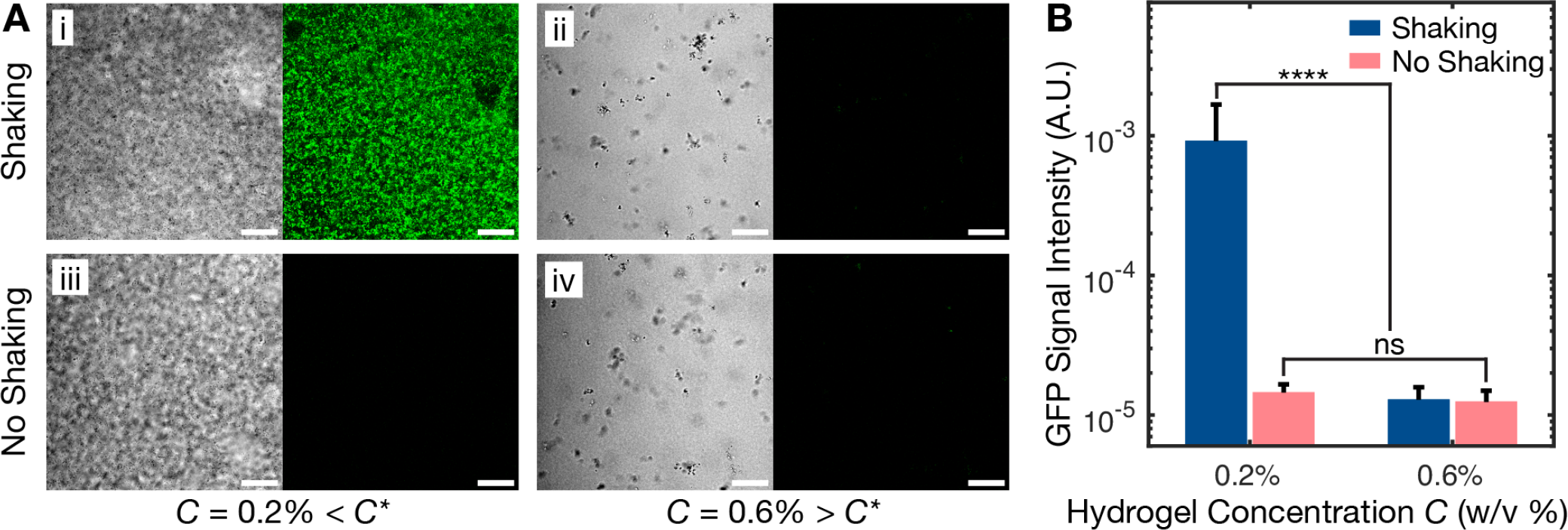
Imaging of oxygen-dependent cellular fluorescence corroborates the hypothesis schematized in Fig. 2. **(A)** Representative confocal brightfield (left) and fluorescence (right) microscopy images of stationary phase, non-motile *E. coli*. Brightfield images show growing cells in all cases; the cells in (i) and (iii) have settled to the bottom, because their matrices are unjammed, whereas the cells in (ii) and (iv) grow in small clusters because the matrix surrounding them is jammed and prevents sedimentation. The cells constitutively express green fluorescent protein (GFP), which requires low levels of oxygen to fold. Only cells grown in shaken aerobic conditions in fluidized matrices with *C < C*^*\**^ have well-folded GFP, and therefore exhibit appreciable fluorescence, as shown in (i). Scale bar = 50 µm. **(B)** GFP fluorescence intensity averaged over planar images taken at a depth 15 µm above the bottom surface of each well averaged over 19-22 replicates confirms that the oxygen-sensitive GFP signal is significantly brighter for cells grown in shaken aerobic conditions in fluidized matrices with *C < C*^*\**^ (leftmost bar). Statistical significance is assessed using a 1-way ANOVA test; **** indicates *p <* 0.0001 and ns indicates a difference that is not significant.

## Results

### Bacteria sharply transition to arrested growth when *C* exceeds *C*^*\**^

We begin by examining the results of standard growth measurements performed in hydrogel-free liquid media in the wells of a 96-well cell culture plate (≈7 mm diameter, ≈11 mm deep) under shaken conditions. As shown by the light pink curve in Fig. 1C, the bacteria follow a complete growth curve with an initial lag phase (0 *< t <* 3 h) followed by exponential growth (3 *< t <* 7h), eventually slowing down and transitioning to stationary phase as they deplete nutrients (*t >* 7h). We observe similar behavior in liquid-like matrices at low hydrogel concentrations (0≤*C* ≤0.3%, pink to orange curves). With progressive increases in *C*, cells slowly begin to transition to stationary phase earlier, yet all cultures reach similar stationary phase OD_600_ values (*t >* 18h)—summarized by the points in the shaded gray region of Fig. 1D. Surprisingly, however, further incrementing *C* slightly from 0.3%(*C < C*^*\**^) to 0.4%(*C > C*^*\**^) results in a sharp transition to dramatically different growth behavior. As shown by the green-blue-purple curves in Fig. 1B, despite the identical initial nutrient conditions and continuous shaking, for all *C* above this transition, cellular growth is slower and consistently arrested at a similarly-low stationary phase cell density. This step-wise transition between continued growth and arrested growth, tuned by a slight change in hydrogel concentration across *C*^*\**^≈ 0.35%, is summarized by the stationary phase OD_600_ values shown in Fig. 1D. We observe the same phenomenon with non-motile *E. coli* (Δ*flhDC*) as well (Fig. S5), suggesting that it is not linked to bacterial motility, but is more general.

What causes this sharp transition from continued to arrested growth across *C*^*\**^? One possibility could be that, as *C* exceeds *C*^*\**^, cells become physically confined in the void space between packed microgels, causing the microgels to deform and exert compressive forces that alter bacterial physiology. Direct mapping of this void space structure in 3D, detailed previously [37], indicates that this is not the case, however. In particular, the characteristic tightest size of these confining voids is larger than the size of a cell body at *C*^*\**^—large enough that motile bacteria can swim through the void space, as we verified independently [36, 37, 58]. Therefore, we expect that the cells are not appreciably compressed by the surrounding microgels at this transition. Moreover, even if slight compression of individual cells were to arise in the experiments, the matrix shear elastic modulus and yield stress at *C*^*\**^ are *∼*1 Pa and *∼* 0.1 Pa, respectively—at least 10^4^ times smaller than the scale of the stresses at which changes in bacterial physiology have been reported to arise [46].

A related possibility could be that, due to the reduced amount of void space available to cells to grow in as *C* increases, crowding of cells causes them to somehow impede each other’s growth. However, at *C*^*\**^, the volume fraction taken up by the swollen microgels is *≈* 64%, which corresponds to random close packing of spheres; thus, *≈* 36% of the overall matrix volume corresponds to liquid-infused void space. At the onset of the transition from continued to arrested growth, the cells only take up ≈0.1% of this void space. Cells are thus not competing for void space at this transition. Indeed, its abrupt nature, induced by just a slight increment of *C* across *C*^*\**^—as opposed to a smooth transition of steadily decreasing growth with increasing *C*, concomitant with the steady decrease in void space volume fraction and pore size—suggests that this transition does not directly arise due to confinement of individual cells. Nor is it associated with changes in the local mechanical environment encountered by individual cells, which also does not abruptly change across *C*^*\**^. We therefore seek an alternate explanation of this phenomenon.

### This growth transition coincides with a sharp transition in matrix rheology

Close comparison of the data in Figs. 1B and D provides a clue to the underlying origin of this phenomenon: the transition between continued and arrested growth occurs when the microgels begin to jam, at which point (*C* = *C*^*\**^) the matrices abruptly transition from being easily fluidized by shaking to tough enough to withstand shaking (detailed in *Materials and Methods*). While the flow field in each microplate well generated by linear plate reader shaking is complex, we can estimate the amplitude of the maximally-imposed shaking stress using the classic solution of Stokes’ second problem [59], 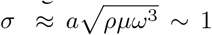, where *a* = 3 mm is the maximal distance traveled during shaking, *ρ* ≈1 g*/*mL is the matrix density, *ω* ≈10 s^*−*1^ is the shaking frequency, and *µ* is the matrix dynamic shear viscosity evaluated at a shear rate equal to *ω* (Fig. S1B). This magnitude of external stress exceeds the highest measured yield stress *σ*_*y*_ ≲ 0.01 Pa for *C < C*^*\**^—hence, these matrices are fluidized by shaking, with the microgels continually rearranging and thereby promoting fluid flow and solute mixing throughout [38]. By contrast, for *C > C*^*\**^, *σ*_*y*_ approaches and then exceeds this value of *σ*, indicating that these matrices remain solid-like upon shaking; the network of inter-grain contacts can support the applied stress, causing individual grains to deform, but not rearrange positions with their neighbors, thereby suppressing interstitial flow and solute mixing [38]. We therefore hypothesize that the transition between continued and arrested growth revealed by the experiments in Fig. 1 reflects a transition in the transport of nutrients throughout the matrices.

### The balance between shaking stress and the matrix yield stress modulates mixing-induced oxygen transport, thereby regulating cellular growth

One likely candidate for a growth-limiting nutrient is oxygen—a metabolite used by *E. coli* to respire. These bacteria are facultative anaerobes: when oxygen is available, the cells can fully breakdown and utilize carbon sources via aerobic respiration, whereas when oxygen is unavailable, the cells only ferment carbon to usable energy, causing growth to become arrested compared to the aerobic case. (For this reason, *E. coli* are traditionally grown in shaken liquid cultures to enhance the transport of dissolved oxygen from the air head space above the culture.) Could oxygen limitations be arresting *E. coli* growth when the hydrogel matrix is solid-like (*C > C*^*\**^), in which case oxygen transport to the cells relies primarily on slow diffusion from the top surface of the matrix? Quantitatively, the dynamics of oxygen concentration 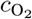 in a static matrix are described by the conservation equation 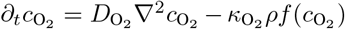, where 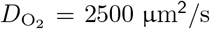 is the oxygen diffusivity, 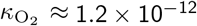 mM (cells*/*mL)^*−*1^s^*−*1^ is the maximal oxygen uptake rate per cell [60], 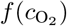 describes the influence of oxygen availability on uptake rate via Michaelis–Menten kinetics taken to be *≈* 1 for simplicity, and we estimate *ρ ≈* 10^9^ cells*/*mL using the maximal value at stationary phase for *C > C*^*\**^. Solving this equation along the one-dimensional coordinate describing the depth into the matrix from its aerated surface then yields an estimate for the characteristic depth to which oxygen diffuses before it is completely taken up by cells [61]: 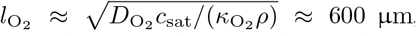, where *c*_sat_ = 180 µM is the saturated level of dissolved oxygen [62]. This depth is *∼*10 times smaller than the matrix height, and the corresponding oxygen depletion time is 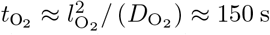, far shorter than the time at which the slowdown and eventual arrest in growth become apparent in our experiments (≈4 h). That is, when the hydrogel matrix is tough enough to withstand shaking (*C > C*^*\**^) and thus oxygen availability is diffusion-limited (lower right schematic in Fig. 2), we expect that ∼90% of the matrix volume rapidly becomes anoxic, causing the cells therein to transition to slower, arrested growth.

This conceptual picture makes a testable prediction: that under non-shaken conditions (*σ* = 0), oxygen availability will always be diffusion-limited, and thus bacterial growth will be similarly slow and arrested even when *C < C*^*\**^. To test this prediction, we repeat the same growth experiments, but without shaking. As expected, in this case, we do not see a sharp transition from continued to arrested growth with increasing *C* across *C*^*\**^; instead, bacteria in all the different matrices show similar slow, arrested growth curves, as shown in Fig. 2A. As summarized by the pink circles in Fig. 2B, all the stationary phase OD_600_ measurements at *C > C*^*\**^ collapse on top of those taken under shaken conditions, shown by the blue circles, indicating that growth is similarly arrested across these hydrogel concentrations.

As another test of this picture, we repeat the same growth experiments, with shaking, but in an anaerobic chamber where oxygen is absent. In this case, we again expect that bacterial growth will be slow and arrested for all *C*; indeed, we expect even slower growth compared to the previous experiments, where oxygen could still diffuse from the top surface of the matrix. Consistent with our expectation, bacteria in all the different matrices of different *C* show similar slow, arrested growth curves, as shown in Fig. 2C. Moreover, as summarized by the yellow circles in Fig. 2B, all the stationary phase OD_600_ measurements are consistent across all values of *C*, and are even lower than those in the experiments conducting in aerobic external settings.

As a final test of our picture, we take advantage of the fact that newly-synthesized GFP requires oxygen to fold properly and exhibit fluorescence [55–57]. Hence, to directly probe cellular growth and oxygen availability in the hydrogel matrices, we repeat both shaking and non-shaking experiments for *C* = 0.2% *> C*^*\**^ and *C* = 0.6% *< C*^*\**^, using non-motile *E. coli* (to eliminate any confounding effects of aerotaxis) that constitutively express GFP. After ∼20 hours of growth, well into the transition into stationary phase, we use confocal microscopy to visualize cell growth and GFP fluorescence at the bottom of each matrix. The results are shown in Fig. 3A. As expected, the only condition exhibiting appreciable GFP signal is that of shaken unjammed matrices (i), for which *σ > σ*_*y*_. The other conditions (ii-iv) all have *σ < σ*_*y*_—jammed matrices are tough enough to be unmixed in both shaken (ii) and non-shaken (iii) conditions, as well as non-shaken unjammed matrices (iv)—and hence have limited oxygen availability, corroborated by the lack of GFP signal. These observations are quantified in Fig. 3B. Taken altogether, these results demonstrate that the balance between the yield stress and external stress regulates bacterial growth by modulating oxygen transport to the cells.

## Discussion

By probing *E. coli* growth inside permeable 3D granular hydrogel matrices, we have shown that the balance between the yield stress *σ*_*y*_ and external stress *σ* can regulate bacterial growth by modulating transport of externally-supplied nutrients. This balance can be quantified using the Bingham number Bi≡*σ*_*y*_*/σ*, a dimensionless parameter used in the field of rheology [63], as summarized in Fig. 4. In particular, our experiments demonstrate that when the cells inhabit strongly-forced (large *σ*) and easily-fluidized (small *σ*_*y*_), and thus well-mixed, environments, nutrients are readily available to the cells and their growth progresses completely (Bi *<* 1 shown in the lower right of Fig. 4). By contrast, when the cells inhabit weakly-forced (small *σ*) and tough (large *σ*_*y*_) environments, slow diffusion from external boundaries limits the availability of growth-limiting nutrients, and cellular growth is arrested (Bi *>* 1 shown in the upper left of Fig. 4. This mechanism by which environmental rheology regulates bacterial physiology by modulating *large-scale* nutrient transport does not arise from *local* mechanical interactions, unlike other mechanisms [41–49, 51–54]; instead, it likely operates in conjunction with them.

**FIG. 4.**
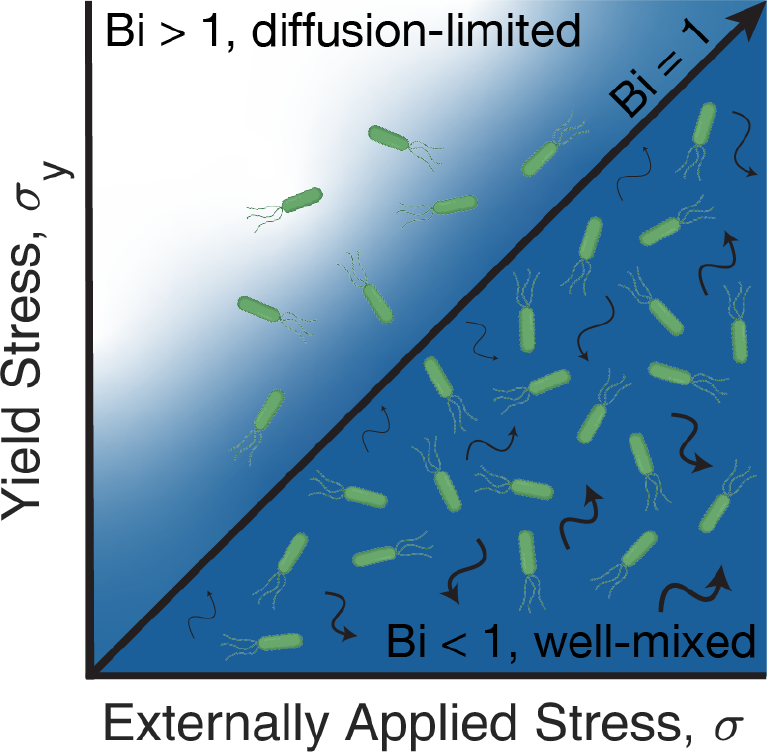
State diagram describing how the balance between environmental yield stress *σ*_*y*_ and the external stress *σ* regulates bacterial growth by modulating nutrient transport. When cells inhabit fragile environments that are fluidized by external forcing (lower right), nutrients (blue) are well-mixed and cellular growth progresses completely. By contrast, when cells inhabit tough environments that are not fluidized (upper left), nutrients are not wellmixed and their availability is diffusion-limited, arresting cellular growth. The boundary between these two growth regimes is described by the diagonal line, Bi≡*σ*_*y*_ */σ* = 1. As shown in Table I, natural bacterial communities inhabit settings that span both regimes of this state diagram.

Our experiments used plate reader shaking to test the influence of forcing, and used tunable granular hydrogel media to test the influence of environmental rheology, enabling us to explore a broad range of Bi in a well-defined and systematic manner. Moreover, as an illustrative example, we used ambient oxygen as a growth-limiting nutrient for cells that grow more efficiently via aerobic respiration. Nevertheless, we expect that our central finding, summarized by Fig. 4, is more generally applicable across different forms of forcing, microbial cell types, and nutrient sources, in diverse environments where nutrients are not uniformly abundant. In fact, given that bacterial habitats (e.g., mucus in the body, extracellular polymer networks in biofilms) have such widely-varying rheological properties and encounter diverse forms of external forcing (e.g., mechanical agitation, imposed fluid flows), Bi spans a broad range of values from much smaller to much larger than 1 in natural and industrial settings—as summarized in Table I. Exploring the generality of our results will therefore be a useful direction for future work.

Moreover, while our study focused on overall bacterial growth, we conjecture that the interplay between environmental yielding and external forcing can regulate other aspects of bacterial physiology as well. For example, we showed that when Bi *>* 1, nutrient availability is limited by the competition between diffusion and uptake by cells. Under these conditions, nutrients are distributed heterogeneously throughout space, potentially driving collective migration and non-uniform spatial organization of cells in a population [64, 65]. Such “patchy” nutrient availability can also promote the establishment of phenotypic and genotypic heterogeneity, as well as competition and cooperation via metabolic cross-feeding, in a population [66]—with implications for e.g., the maintenance of genetic diversity in the population, as well as its resilience to external stressors such as administered antibiotics [67–70]. Investigating how our findings may translate to other such changes in bacterial physiology will be an interesting avenue for future research.

## Acknowledgements

This work was supported in part by the National Science Foundation Graduate Research Fellowship Program (to A.M.H.) under grant no. DGE-2039656, Princeton University’s Materials Research Science and Engineering Center under NSF grant no. DMR-2011750, and partial support from NSF grants CBET-1941716, DMR-2011750, and EF-2124863. Any opinions, findings and conclusions or recommendations expressed in this material are those of the authors and do not necessarily reflect the views of the National Science Foundation. This work was also supported in part by a Camille Dreyfus Teacher-Scholar Award from the Camille and Henry Dreyfus Foundation, the Pew Biomedical Scholars Program, and the Princeton Catalysis Initiative. We thank Christopher Browne, Jenna Moore-Ott, and Victoria Muir for useful discussions; the lab of Mohamed Donia for allowing use of the anaerobic chamber; and the lab of Howard Stone for allowing use of the roughened parallel plates for rheology.

## Author contributions

A.M.H. and S.S.D. designed the experiments; A.M.H. performed all experiments; A.M.H. and S.S.D. designed and performed theoretical calculations; A.M.H. and S.S.D. analyzed all data, discussed the results and implications, and wrote the manuscript; S.S.D. designed and supervised the overall project.

---

**FIG. S1.**
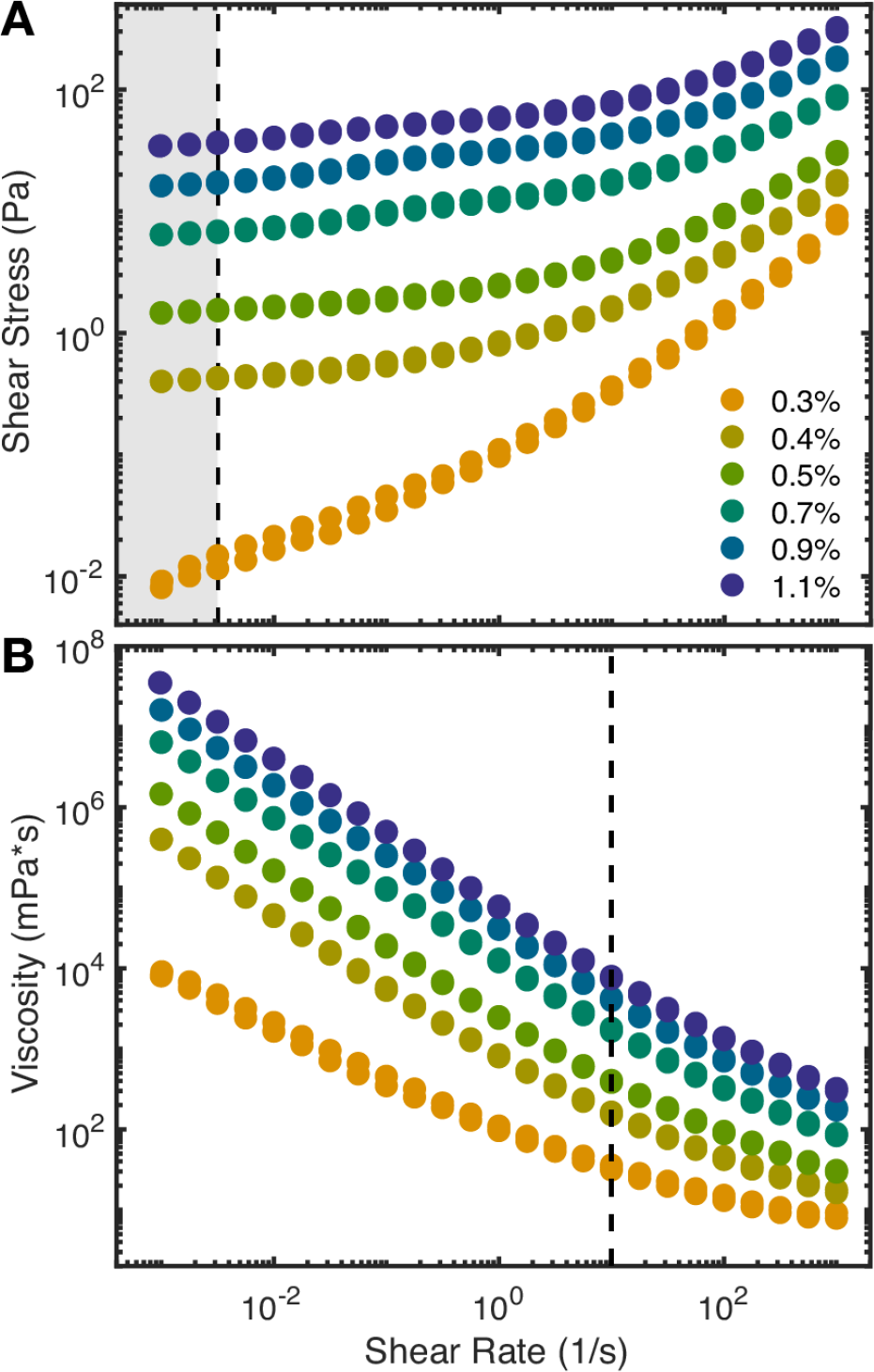
Unidirectional steady shear rheology measurements of granular hydrogel matrices. **(A)** Measurements of shear stress *σ* as a function of applied unidirectional shear rate 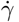 . For sufficiently large *σ > σ*_*y*_, each matrix is fluidized and *σ* increases with increasing 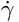 ; the corresponding slope, which yields the dynamic shear viscosity, is plotted in **(B)**. Extrapolating that shear stress measurements to 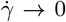 by averaging the measured shear stress for 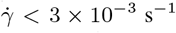 (shaded grey region) then provides an estimate for *σ*_*y*_ . For visual reference, the dashed line in **(B)** indicates the estimate maximal shear rate (10 s^*−*1^) encountered during linear plate reader shaking. The legend indicates the dry mass fraction of granular hydrogel used to prepare the matrices.

**FIG. S2.**
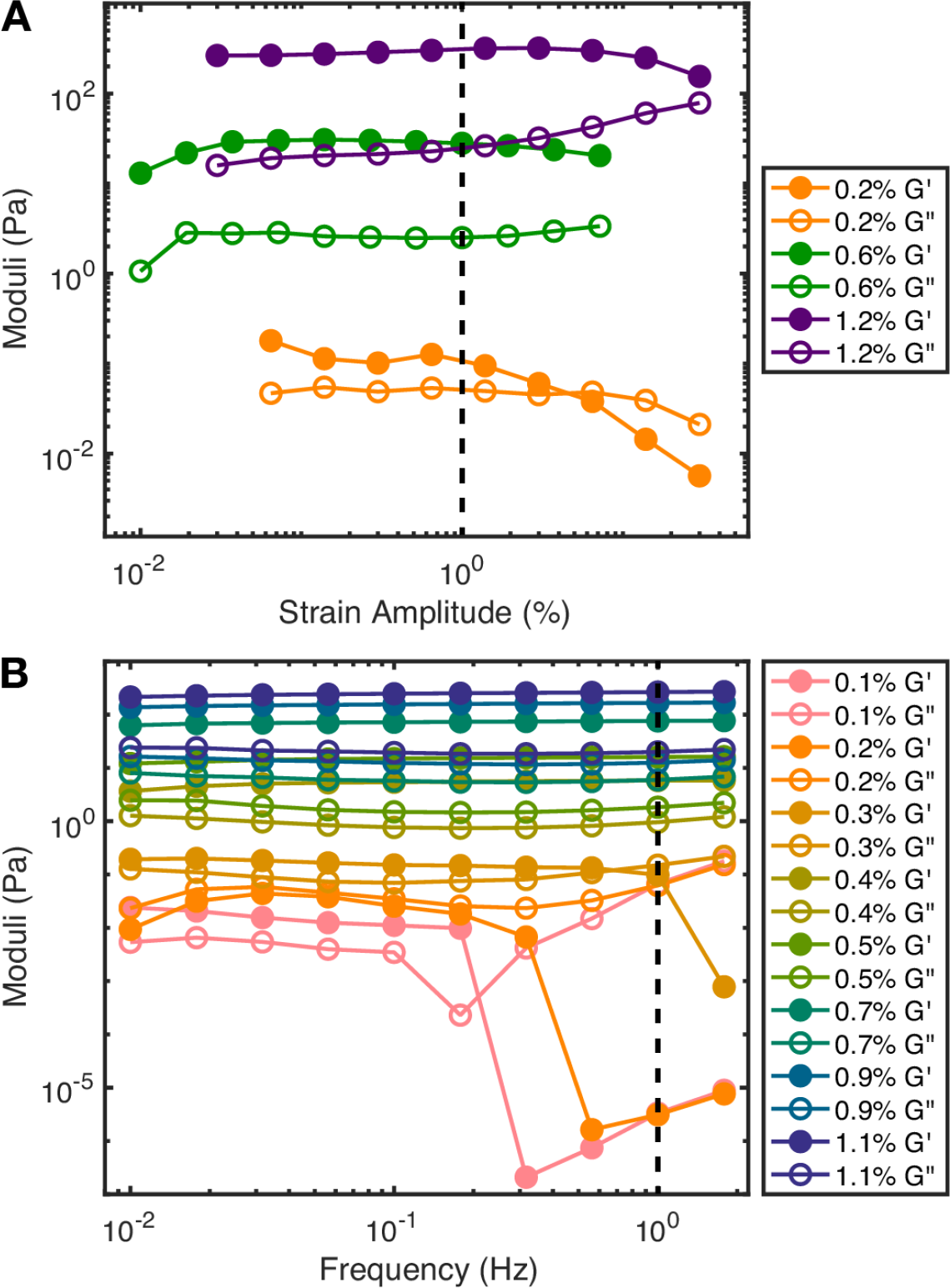
Oscillatory shear rheology measurements of granular hydrogel matrices. Measurements of the elastic storage (closed circles) and viscous loss (open circles) shear moduli, *G*^*′*^ and *G*^*″*^ respectively, as a function of **(A)** strain amplitude at a fixed oscillation frequency of 1 Hz, or **(B)** oscillation frequency at a small fixed strain amplitude of 1%. The legend indicates the dry mass fraction of granular hydrogel used to prepare the matrices.

**FIG. S3.**
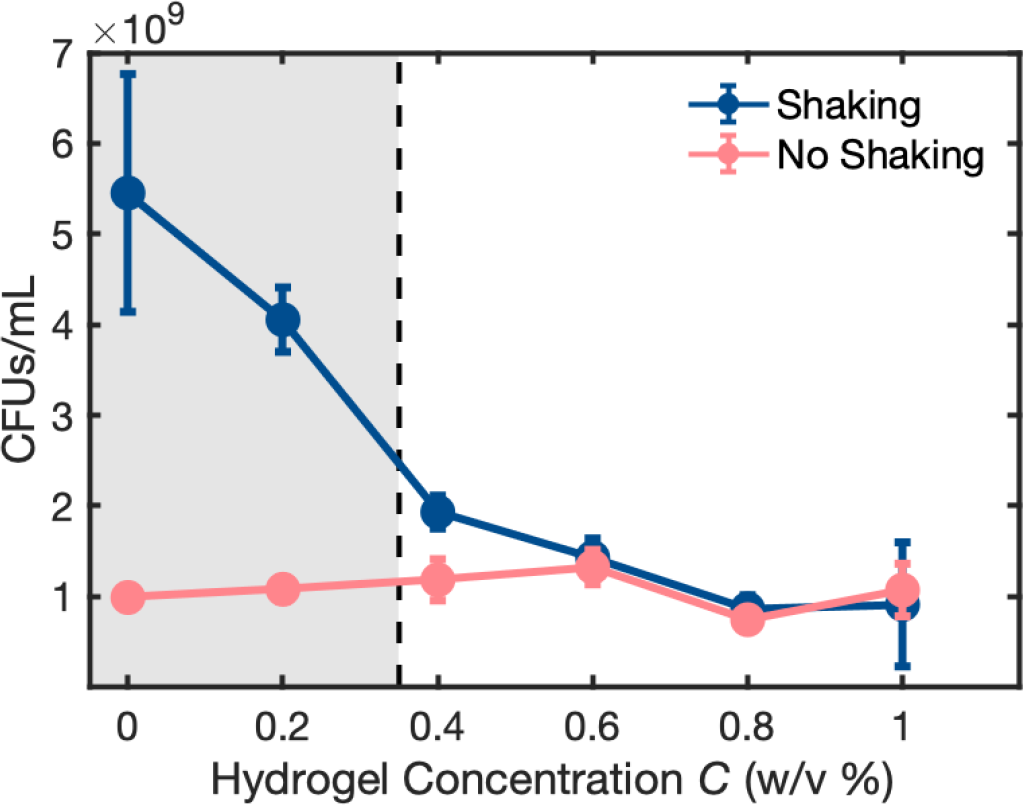
Cell viability measured by colony forming units of cells grown in shaken and non-shaken conditions for 24 h. The data show a similar trend to the results describe in the main text, which were obtained using OD_600_ measurements instead. Dashed line marks the hydrogel jamming concentration. Error bars represent standard deviation among quadruplicates from 1 experiment.

**FIG. S4.**
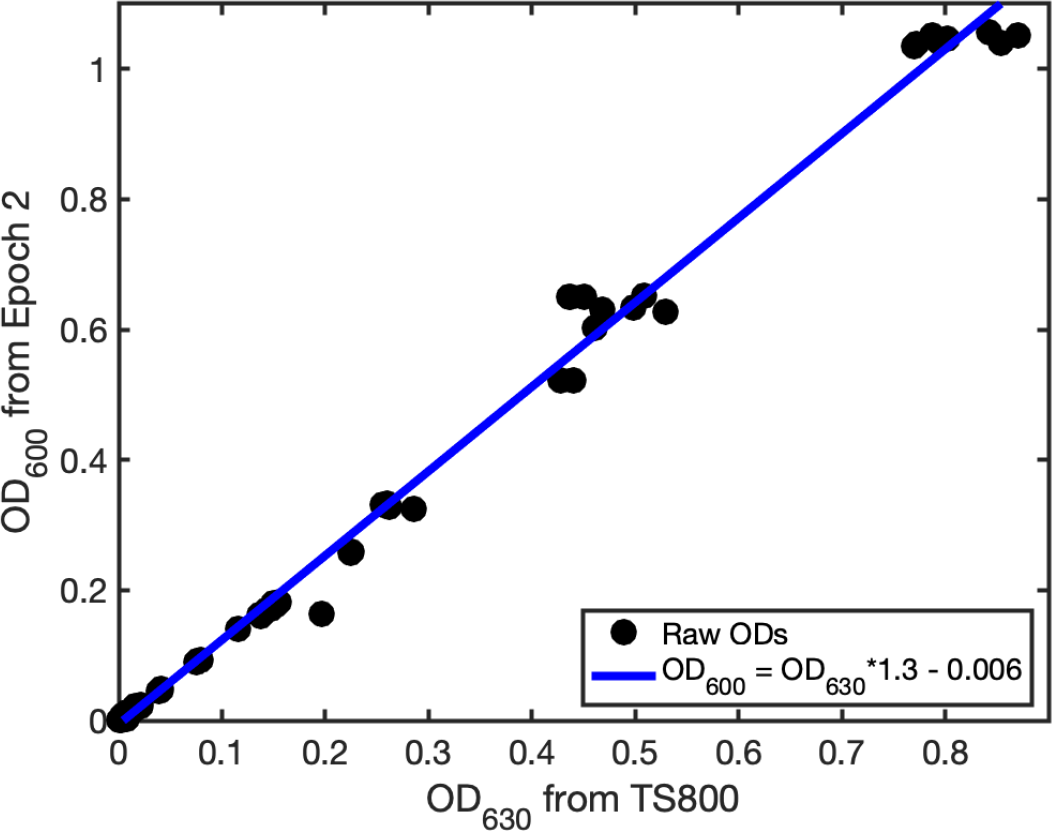
Conversion between OD_630_ and OD_600_ measurements. We measure the OD of 96 well plates containing the same dilutions of stationary phase *E. coli* in the Biotek Epoch 2 (OD_600_) and Biotek 800TS (OD_630_) plate readers. The best fit line shown in blue is used to convert between OD_630_ and OD_600_ measurements.

**FIG. S5.**
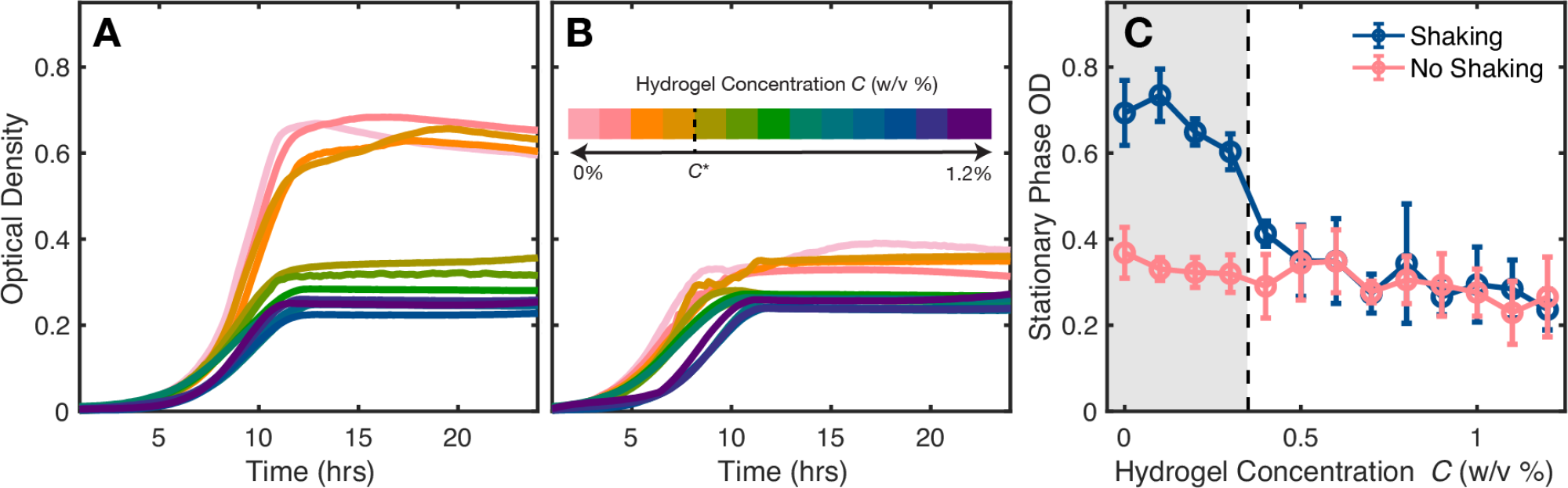
Non-motile *E. coli* show similar growth behaviors as the motile strains described in the main text figures. Average growth curves of non-motile *E. coli* in varying concentrations of granular hydrogel measured with spectrophotometry under **(A)** shaken and **(B)** non-shaken aerobic conditions. For **(A)** and **(B)**, each curve is an average of 3-5 replicates from 1 experiment. **(C)** Stationary phase optical density, defined as the average OD_600_ from *t* = 18*−*24h, for shaken (blue) and non-shaken (pink) growth curves. For shaken conditions, stationary phase OD_600_ drops at a threshold concentration that matches the hydrogel jamming concentration *C*^*\**^. For *C > C*^*\**^, shaken and non-shaken stationary phase OD_600_ measurements collapse, just as in the case of motile *E. coli* as shown in the main text. Dashed line marks the hydrogel jamming concentration. Error bars represent standard deviation from replicates across 2-4 independent experiments.

